# Diffusional variance-determined stochastic translation efficiency

**DOI:** 10.1101/2022.03.13.483370

**Authors:** Attila Horvath, Yoshika Janapala, Ross D. Hannan, Eduardo Eyras, Thomas Preiss, Nikolay E. Shirokikh

## Abstract

Full-transcriptome methods have brought versatile power to protein biosynthesis research, but remain difficult to apply for the quantification of absolute protein synthesis rates. Here we propose and, using modified translation complex profiling, confirm co-localisation of ribosomes on messenger(m)RNA resulting from their diffusional dynamics. We demonstrate that these stochastically co-localised ribosomes are linked with the translation initiation rate and provide a robust variable to model and quantify specific protein output from mRNA.

## MAIN

Translation of mRNA into proteins is an actively regulated stage of gene expression^1,2^. Many of the rapid responses of cells are solely based on or are induced by translational control, including in prominent and archetypal biological processes such as nutrient-induced regulation, unfolded protein stress response, dynamic cell reprogramming in embryonic development and synaptic plasticity^2–7^. Yet, it remains a substantial challenge to accurately determine translation rates transcriptome-wide^8,9^.

A breakthrough in translation investigation methods has been offered by combining ribonuclease (RNase) protection approach (generation of ribosomal footprints on mRNA by digesting away the unprotected mRNA parts with RNase) and high-throughput RNA sequencing, in the form of ribosome and translation complex profiling (TCP-seq), and a range of derived methods (**Figure 1a**)^8,10,11^.

**Figure 1.**
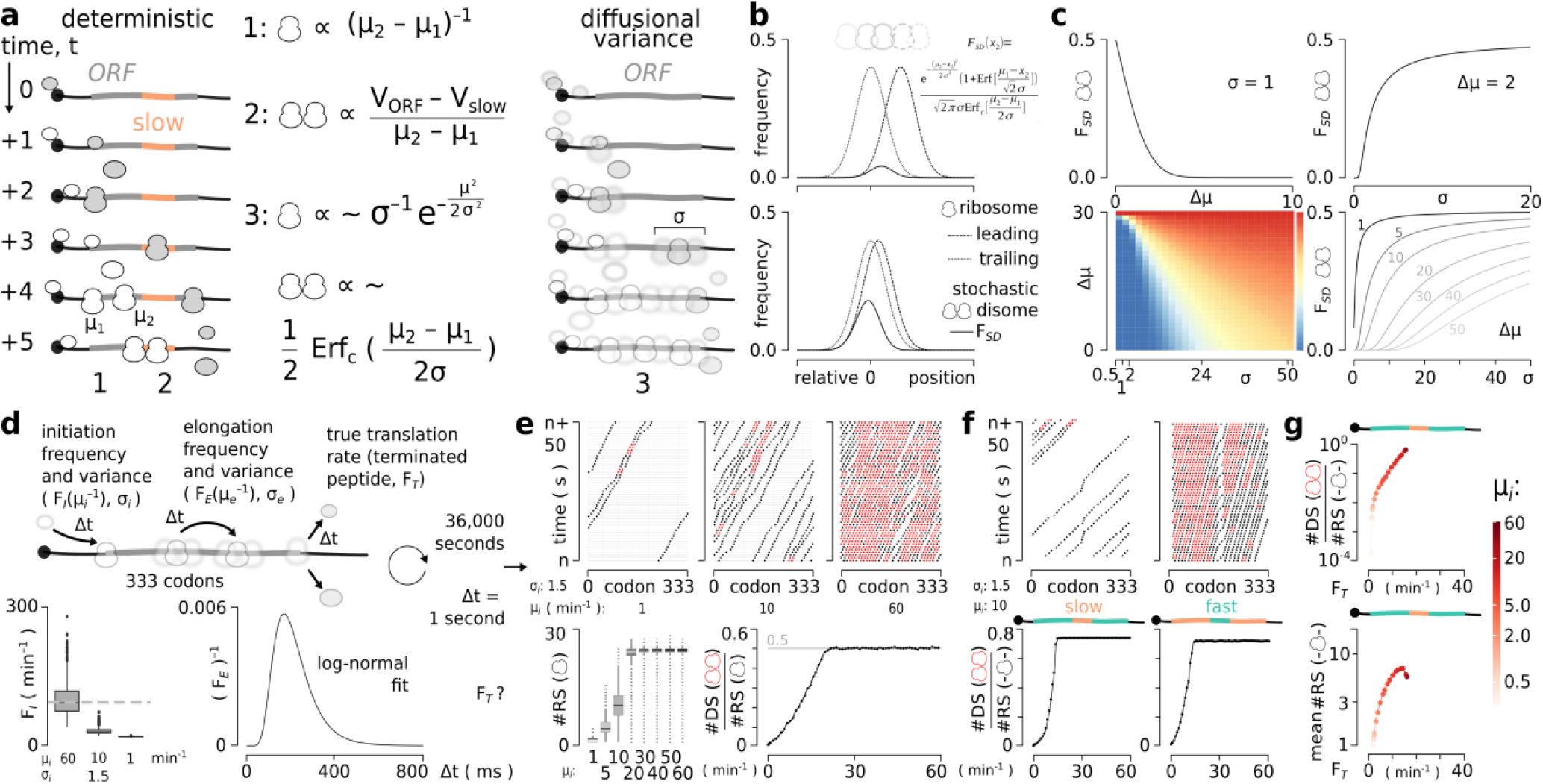
Stochastic co-localised ribosomes (disomes, DS) can result from diffusional initiation and elongation rate variances and are monotonously linked to the translation initiation rate. **(a)** Comparison of the classical ‘deterministic’ view (*left*) on the ribosomal (RS) progression on mRNA with a diffusion-aware stochastic model (*right*). Note that in the classical model, DS can only occur at certain sites with slow elongation rate or arrested translation. **(b)** Generalised mathematical model of stochastic disome occurrence (stochastic disome frequency, F*_SD_*) for an infinitely long model mRNA containing two RS (see more in ‘Mathematical modelling’ in Methods). **(c)** F*_SD_* for case as in (b) monotonously depends on initiation frequency across all variance values. **(d)** Computational diffusion-aware model of mRNA translation accounting for the specific experimentally-determined distributions of the initiation and elongation rates and variances (frequency of initiation F_*I*_ and frequency of elongation F_*E*_, respectively see more in ‘Construction of *in silico* translatome and simulation of translation supported by *in vivo*-derived parameters’ in Methods). **(e)** Model from (d) realised across different median initiation rates and using a homogeneous F_*E*_ log-normal distribution of elongation dynamics for the entire simulated Open Reading Frame (ORF) of 333 codons. Temporal (Y-axis) traces of individual RS positions on mRNA (‘trace plots’) for slow, medium and fast initiation rate cases are shown across the consecutive mRNA codons (X-axis) (*top*). Note that even a slow initiation rate with sparsely situated RS results in stochastic co-localisations. Steady-state number of RS over the model 333-codon ORF is dependent on the initiation rate, until saturation is reached (*bottom left*). DS to RS ratio at the steady state unambiguously depends on the initiation frequency (*bottom right*). **(f)** Model realisations as in (e) with the initiation rate sampled from the range of initiation for mRNAs with heterogeneous F_*E*_ with a first ‘fast’ segment (sampled from the upper quartile of the elongation rate distribution) (*left*), a 21-nucleotide-long ‘slow’ segment (sampled from the lower quartile of the elongation rate distribution) in the middle representing a ‘stalling’ or slowdown site, and a third ‘fast’ segment (sampled from the higher quartile of the elongation rate distribution) (*right*), accompanied with the inverse model ORFs. **(g)** Translation rate (protein yield as determined by the terminated peptide frequency F*_T_*) retains monotonous dependency on DS to RS ratio, as opposed to the mean monosome (MS) density on mRNA (equivalent of the normalisation of RS to RNA signals in experimental profiling data).

In this study, we identify a new type of signal that can be observed within translation complex profile sequencing data generated in experiments that involve rapid crosslinking of *in vivo* translational complexes^12–15^. The signal originates from the stochastic co-localisation of individual ribosomes on mRNA, and is evidenced by long disome-derived (possibly but rarely, multisome-derived) footprints located outside the regions of non-random elongation stalls (**Figure 1a**).

Many of the existing works have characterised ‘collided ribosomes’^16–25^, yet nuclease-resistant disome occurrence has been mostly attributed to discontinuous elongation rates, such as in ribosomal stalls or slowdowns^17–19,24^. Attempting to generalise this problem, we asked what could be a minimal prerequisite of ribosomal co-localisation on mRNA (**Figure 1a**). Using a theoretical mRNA with infinite ORF length and featuring only two ribosomes, in our model a co-localisation takes place when the trailing ribosome would attempt to elongate further on the message than the leading ribosome. Consequently, unlike in traditional representation where ribosomal positions are deterministically defined, we used a more realistic assumption that diffusional heterogeneities of the elongation rate create a probabilistic function of ribosome localisation, if two or more ‘independent’ ribosomes are considered^26–28^. Simplifying this distribution to normal and identical in standard deviation for all ribosomes (with μ representing distance between ribosomes over mRNA defined as the inverse initiation frequency), this model results in a normal-like stochastic disome occurrence (**Figure 1b**). Intuitively, disome probability maximum does not exceed 0.5 for two ribosomes. The disome probability also non-linearly but monotonously increases depending on the ribosome density (*i.e*. initiation frequency), and is dependent on the localisation variance (σ), which cumulatively embodies diffusional variances of all contributing processes (**Figure 1c**).

To attest the potential of theoretical stochastic co-occurrence of the ribosomes in a lifelike scenario, we constructed a parametrised model somewhat inspired by the classical solutions to the road traffic, M/M/1-queues and totally asymmetric simple exclusion process (TASEP; and its derivatives) model of protein synthesis (**Figure 1d**)^29–33^. We did not wish to use this model to investigate finer kinetic coefficients or sub-steps, and rather focused on a solution that would find its base in experimentally measurable variables, with a minimal number of assumptions. Briefly, we first allowed ribosomes to attach to mRNA (‘initiate’) containing a finite long ORF (999 nucleotides) with a certain frequency distribution modelled after *in vivo*-measured initiation parameters^34^, and then investigated step-wise relocation of the ribosomes downstream (‘elongation’), using the reconstructed probability distribution of *in vivo*-measured elongation rate and variance^35^. We then cycled the model for 36,000 steps (model seconds), noting the co-localisation of two ribosomes (formation of disomes; **Figure 1e**). As a result, disomes could be detected in all cases (including those of fully homogeneous, slow or fast elongation rates; **Figure 1f**), and in the cases of discontinuous elongation rates they formed areas of differential frequency of occurrence. We then recorded true protein output by evaluating 378 simulations across a spectrum of initiation rates and codon patterns, and measuring the number of ribosomes translocated from the ORF end as the frequency of termination (F*_T_* or true protein output). The most distinctive, and somewhat unexpected, feature of the model has been the monotonous and largely linear response of the disome to ribosome ratio to the true protein output, across all ORF types (**Figure 1g**). Critically, this ratio is a result of the self-normalisation of two values of a similar nature, which can remove biases deriving from methodological and instrumental differences and, importantly, provide a capability to directly quantify absolute protein biosynthesis rates.

Having proven that stochastic disomes could be, in principle, observed within the known initiation and elongation variances and rates, we employed a specialised version of translation complex profile sequencing (TCP-seq)^12-14,36^ in an attempt to identify them *in vivo*. We used rapidly-crosslinked translational complexes separated into total RNA (*T*), total translated RNA (*tt*), polysome-engaged ribosomal small subunit (SSU), ribosome (RS) and disome (DS) fractions (**Figure 2a**). Our rigorous bioinformatics analysis revealed that the DS fraction prominently demonstrated the presence of longer footprints (**Figure 2a**), whereas SSU and ribosome fractions mapped in accordance to the previously-published results^12,14^. TCP-seq disomes produced a complex pattern of footprint lengths (**Figure 2a**). We were able to identify the presence of ‘long’ (~64 nt) and ‘short’ (~32 nt) disome footprints, as well as a few intermediate fractions, likely produced by all four ribosomal combinations (with open A-site/’rotated’ giving short and closed A-site giving long footprints) in a disome. We also detected short (‘monosomal’) footprints reflecting these states, as before. While some of the ‘monosomal’ footprints in the disome fraction may result from the cleavage of mRNA between the ORF-neighbouring ribosomes (as these remain associated through an mRNA-independent crosslink), these footprints can also derive from the crosslinking of ORF-distant, singular ribosomes, without a requirement of their tight serial packing on mRNA (**Figure 2b**). Such short footprints in the disome fraction can thus relate to the sites of stable spatial proximity in the higher-order (polysomal) structures, providing an entirely new dimension to the footprint data with a potential for novel insights into translational and spatial dynamics of polysomes, but remain outside the scope of this work.

**Figure 2.**
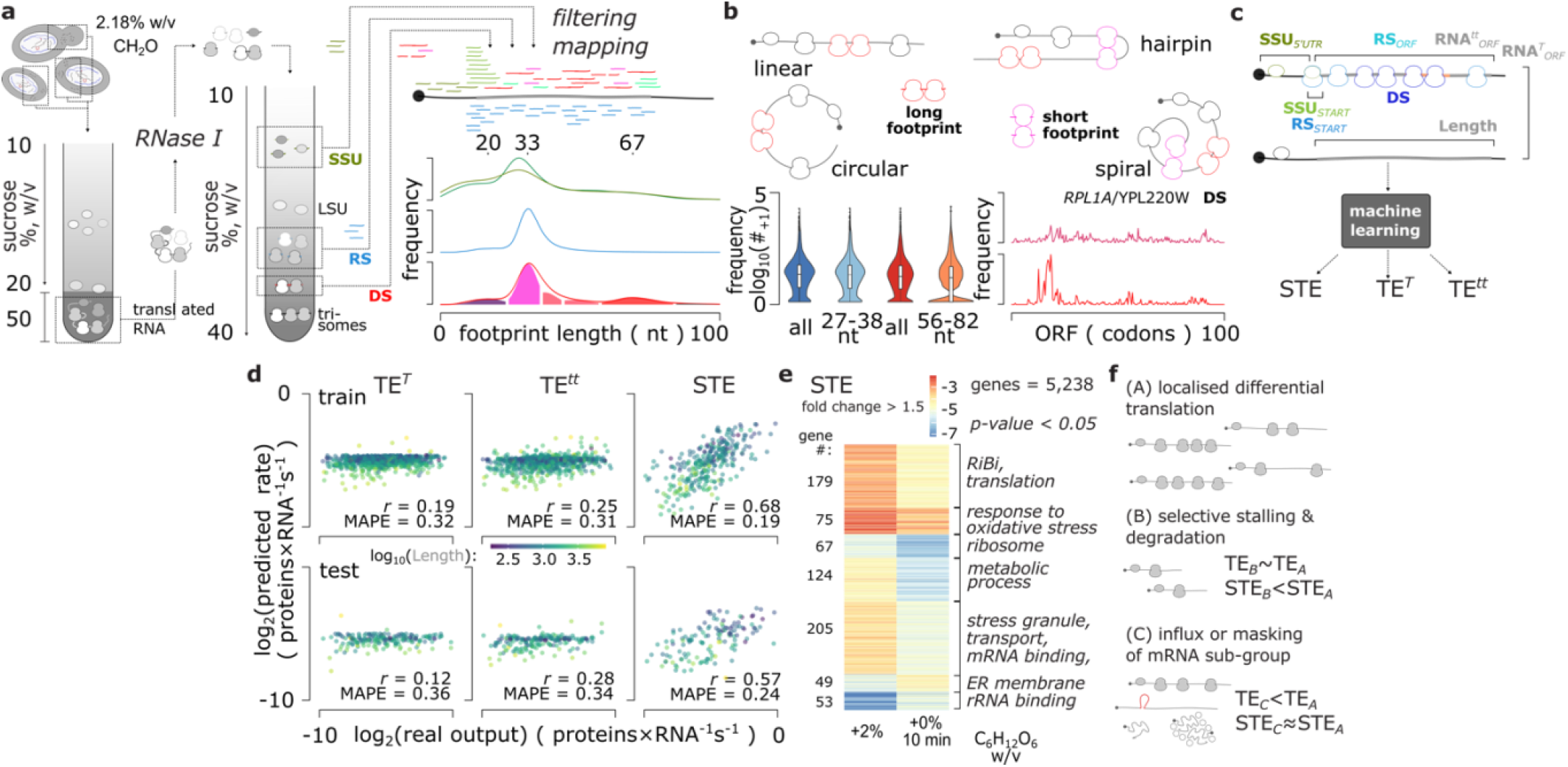
Stochastic Translation Efficiency (STE) measure maps to the experimentally-determined absolute protein synthesis rates and dissects classes of mRNAs with different power in translation. **(a)** Outline of an extended TCP-seq approach used to perform STE calculations. Schematic illustrating the sedimentation-based separation of total translated RNA (RNA^*tt*^), and its RNase I cleavage and fractionation, to generate SSU, RS and DS footprints (*left*). Footprint mapping from different fractions and the respective fragment length distributions (*right*). Note the presence of an additional peak with ~67 nt mode in the DS fraction. **(b)** Features and theorised origins of the DS footprints. Different types of commonly observed polysome arrangements can facilitate contacts between distal RS, resulting in localised ‘short’ footprints associated with the DS fraction, in addition to the ‘long’ DS footprints as a result of stochastic or non-random proximal co-localisation, and ‘short’ footprints due to their partial cleavage in halves (*top*). Many transcripts have RS (blue) and DS (red) fractions with substantial frequency of the footprints in the 27-38 nt (‘short’) and 56-82 nt (‘long’) characteristic footprint ranges, respectively (*bottom left*). DS footprints filtered into the ‘short’ (purple, top) and ‘long’ (red, bottom) groups for a well-covered *RLP1A* mRNA ORF demonstrate partially concordant coverage, with several peaks appearing as unique to each of the groups (*bottom middle*). **(c)** Machine learning approach featuring separate training and cross-validation on two, non-overlapping subsets of the training data. The features used to quantify STE are like-to-like normalised values, as opposed to the Translation Efficiency-like TE^*T*^ and TE^*tt*^ calculated by normalising RS ORF signal to Total (*T*) or total translated (*tt*) RNA-seq signals, respectively. Note all models include ORF length as an input. **(d)** STE demonstrates high correlation with the experimentally-defined protein synthesis rates (metabolic labelling-based quantitative mass spectrometry data from [32]). **(e)** Classes of translationally up- or down-regulated mRNAs during yeast response to the 10-minute glucose depletion as determined by STE. The classes were inferred using k-means clustering of mRNAs with STE fold-change of at least 1.5 between the non-starved and starved states. **(f)** Schematic illustrating robustness of the STE measure of translational power in the background of rapid transcription-, degradation- or masking- and phase separation-induced alterations of the mRNA abundance and availability to translation, allowing to isolate mRNA-specific, UTR-driven translational control *in vivo*.

We next sought to identify the stochastic component of the disome footprints, to use it for protein output modelling as theorised above. We first isolated ‘long’ (56-81 nt) footprints for the disomal and ‘short’ (27-38 nt) footprints for the ribosomal fractions. As the long disome footprints are virtually absent in the ribosomal data (**Figure 2a**), the remaining signal directly supports the stochastic appearance of ribosomal co-incidence and accidental tight packing (**Figure 2b**). Upon all filtering, we inferred the disome to ribosome (DS/RS) signal, and compared against the known accurate measurements of protein synthesis rates in *S. cerevisiae* performed using metabolic labelling and mass-spectrometry^37^. Filtered DS/RS values returned strong positive correlation to the protein output for the mRNAs with a robust stochastic disome signal, directly confirming our theoretical conclusions (data not shown).

We then employed a machine learning approach (**Figure 2c**) which used inputs from ORF length-normalised footprint densities of different fractions, and various measures of translation including DS to RS ratios, initiation and elongation efficiencies, and the ORF length. Upon training, our model returned ~0.6-0.7 correlation with the mass-spectrometry-based measurements benchmarked with an independent data subset not used in the training process (30% of the whole data set). Surprisingly, we could not achieve a meaningfully positive correlation when using ribosomal footprint counts normalised to either total or total translated RNA-seq signals (TE^*T*^, TE^*tt*^; **Figure 2d**). Remarkably, the inclusion of ORF length demonstrated only moderate improvements. Because the DS/RS diffusion variance-derived component is unique to our modelling, we deemed our measure as ‘Stochastic Translation Efficiency’, or STE. Importantly, STE provides a direct estimate of the absolute protein output per mRNA per unit of time, and thus can be used to rank mRNAs and respective UTRs by their power to compete for the initiating ribosomes at any given condition.

Applying STE to a realistic scenario of nutrient starvation (10-minute glucose depletion) in *S. cerevisiae,* we could observe gene clustering by response type and magnitude and reveal finer details of gene expression control. In the non-starved cells, 1,310 genes appeared in the top 25% and 1,310 genes – in the bottom 25% of STE values, with maximal F_*T*_ values approaching 0.055 p/s (**Figure 2e**). Ribosome biosynthesis (RiBi), sugar and amino acid metabolism and biosynthesis of secondary metabolites transcripts were enriched in high STE, whereas endocytosis appeared translationally suppressed (**Figure 2e**). In the starved cells, RiBi transcripts were translated less efficiently (variably), amino acid and secondary metabolism transport and stress granule-related transcripts substantially inhibited, while endoplasmic reticulum (ER) membrane-related mRNAs accelerated their translation (**Figure 2e**). Notably, *DFM* / YDR411C (endoplasmic reticulum (ER)-localised protein; involved in ER-associated protein degradation (ERAD), ER stress, and homeostasis), *OSM1* / YJR051W (*f*umarate reductase, catalyses the reduction of fumarate to succinate; required for the re-oxidation of intracellular NADH under anaerobic conditions; acts as electron acceptor in mitochondrial inter membrane space; has two translation start sites, one at the annotated start codon which produces an ER-targeted form required for anaerobic growth), *RPS21A* / YKR057W (SSU protein), *TPI1* / YDR050C (triose phosphate isomerase, abundant glycolytic enzyme involved in the glycolytic breakdown of carbohydrates into pyruvate; localises to mitochondria and plasma membranes), *PGK1* / YCR012W (phosphoglycerate kinase involved in glycolysis and gluconeogenesis; localises to the plasma membrane, mitochondria, and cytoplasm) were highly translationally up-regulated, underpinning the importance of cytoplasmic gene control in nutrient response. An entirely unexpected finding has been that in the glucose-starved cells, the majority of transcripts demonstrated higher translation rates compared to the non-starved cells (medians of ~0.0052 *vs*. 0.0049 p/s, respectively). This stress-induced translational acceleration may be a consequence of a lesser competition for translation components as most of mRNA becomes unavailable to translation in the starved cells (as also revealed by an excess of monosomes)^36^. Stressed cells also demonstrated high degree of selective translation control, with 645 genes swapping the top and bottom quartiles of the STE. Notably, a signature of accelerated RNA restructuring was observed, where RNA metabolism genes (rRNA maturation, transcription and gene expression) were up-regulated, whereas general translation, nucleotide and ribose phosphate/carboxylic acid metabolism were down-regulated (**Figure 2e**).

To summarise, our work has identified new types of co-localised ribosomes on mRNA, stochastic and spatial disomes, in addition to the commonly-accepted collided ribosome disomes resulting from stalling or local elongation slow-downs. Footprints form the newly-identified types of co-localised ribosomes carry information about single-molecule events such as diffusion- and spatial configuration-driven transient contacts between ribosomes, as they form polysomes with translated mRNA *in vivo*. The stochastic disome signal, together with the other measurements derived from TCP-seq, provide a link to the absolute translation initiation and protein biosynthesis rates. Using STE, it is possible to rank mRNAs by the absolute protein output and thus, characterise the ‘power’ of translation control elements across transcripts in a single setting or between different conditions. STE does not use bias-inducing normalisation to the RNA abundance or signals of different types and relies on self-normalised signal pairs. STE thus evades imprecisions arising from the different accessibility across mRNAs to translation machinery (such as of mRNA located in nucleus, condensed/phase-separated foci *etc*.) and coincidental transcription- and stability-driven mRNA abundance alterations (**Figure 2f**). With STE, we demonstrate that the prototypical example of translational control during yeast response to glucose depletion is more complex than previously thought and includes translational acceleration as well as re-adjustment of the RNA metabolism to suit new translational demands. More details can be uncovered in the future, as STE provides a robust measure permissive to finer dynamics dissection and elucidation of very rapid cell responses suitable for investigating protein biosynthesis over a rapidly changing transcriptome background as occurs in cell stress response and reprogramming.

## MATERIALS AND METHODS

### Mathematical modelling

The developed mathematical model of disome formation was verified by generating positions of ribosomes over 10 million iteration. Then the individual ribosome position was modelled using Gaussian distributions with identical standard deviations (σ = 1) and their relation was assessed in R using the stats package. The resulted probabilities and distributions were also verified in Mathematica^38^. Line and boxplots were plotted using the ggplot2 R package.

### Published data used in the study

To generate a spectrum of elongation rates over codons, a ribo-seq-derived data was used^39^. Initiation rate distribution was reconstructed using by a maximum frequency-limited log-normal distribution fitted to available experimental data^34^. To train and benchmark the developed ensemble models, protein synthesis rate was taken from a study that employed metabolic labelling and mass-spectrometry^32^.

### Construction of *in silico* translatome and simulation of translation supported by *in vivo*-derived parameters

To generate the distribution of elongation rates over the triplets the ribo-seq-derived data^39^ was fitted by log-normal distribution. A log-normal distribution modelling was also employed for initiation rate reconstruction using available experimental data^34^. Maximum values for the initiation rate (number of initiating ribosomes per unit time) were restricted to 60 initiation events per minute for each individual model mRNA. The *in-silico* translatomes were modelled with a prototypical ORF consisting of 333 codons, using the fitted elongation speed profile. Simulations were run for 36,000 model seconds (10 model hours) with a range of fitted initiation (μ = 1, 10, 20, 30, 40, 50, 60) and elongation rate distributions. ‘Slow’ and ‘fast’ segments were taken from the lower and upper quartiles of the elongation rate distribution, respectively, while the trimodal patterns of fast-slow-fast and slow-fast-slow were constructed with a middle segment of 7 codons (21 nt) surrounded by either a fast or a slow segment. Line, trace and box plots were generated using the R package ggplot2.

### Cell material, fixation regimen and cytosol collection

Cell fixation and harvesting was generally performed as described before^40,41^ with modifications^36^. Wild-type (WT) yeast of cell line BY4741 (MATa his3Δ1 leu2Δ0 met15Δ0 ura3Δ0) was grown in 1 litre of YPD (yeast extract (Merck/Sigma-Aldrich cat. no. 70161), peptone (Merck/Sigma-Aldrich cat. no. 70178), 2% dextrose (Merck/Sigma-Aldrich cat. no. 49139), 40 mg/L adenine sulphate (Amresco 0607-100G) media till they reached an optical density of 0.7-0.8 AU at 600 nm (OD600). The cells were immediately snap-chilled by mixing with 25% w/v of crushed ice and 37% formaldehyde (solution (methanol-stabilised solution; ‘formalin’; Merck/Sigma-Aldrich cat. no. F11635-500ML) was added to a final concentration of 2.2% w/v immediately under constant mixing. Cells were incubated for 10 min on ice for fixation and then pelleted by centrifugation at 4°C, 5,000×g for 5 min. The fixed cell pellet was resuspneded and washed with 40 ml of buffer A (20 mM HEPES-KOH pH 7.4 at 25°C, 100 mM KCl and 2 mM MgCl2), followed by centrifugation at 4°C, 5,000×g for 5 min. The supernatant was discarded and the fixed and washed cell pellet was resuspended in 40 ml of buffer A1 (20 mM HEPES-KOH pH 7.4 at 25°C, 100 mM KCl, 2 mM MgCl2 and 250 mM glycine). This wash is critical for avoiding irreproducible crosslinking and the buffer addition must not exceed 20 min of the total time from the beginning of the cells’ harvest. The cells were pelleted again by centrifugation at 4°C, 5,000×g for 5 min, and the buffer A1 wash was repeated two more times. The washed cell pellet was aspirated (~1 g wet cell mass) and resuspended in 550 μl of buffer A2 (buffer A1 supplemented with 5 mM DTT, 1 U/μl RNaseOUT RNase inhibitor (Thermo Fisher Scientific) and 1× Complete EDTA-free Mini Protease Inhibitor (Merck). The cell suspension was then flash-frozen by dripping into liquid nitrogen and the frozen suspension pellets stored at-80°C.

To disrupt the cell wall and membrane, a 10 ml stainless steel grinding jar (Retsch) was pre-cooled by partial submerging in liquid nitrogen and then filled with ~2 g of the frozen cell suspension pellets and two 12 mm stainless steel grinding balls (Retsch). The sealed grinding jars were shaken at 27 Hz for 1 min in MM400 mixer mill (Retsch), re-cooled by partial submerging in liquid nitrogen and continued shaking for 1 min more. The resultant powdered grindate was stored in 1.5 ml low protein binding tubes (Eppendorf) at −80°C in ~100 mg aliquots and used as necessary. Usually, 600 mg of the grindate was used per one experiment comprising polysome sedimentation profile analysis, separation of the cytosol into translated and non-translated fractions, and further separation of the translated fraction into the ribosomal small subunit (SSU), ribosome (RS) and disome (DS) fractions upon RNase digestion.

### Separation of the fixed (poly)ribosomal complexes away from the non-translated fractions of the cytosolic RNA

We generally followed the procedure established by us earlier^40,41^ to enrich for ‘total translated’ RNA (*tt*) based on its co-sedimentation with (poly)ribosomes, but introduced a more refined approach^36^. A modification has been made to allow for a direct separation monitoring using absorbance profile readout upon ultracentrifugation and dismissal of the re-solibilisation step that resulted in higher material losses and excessive denaturation and aggregation previously. ~100 mg of the frozen cell grindate were thawed, supplemented with 150 μl buffer A2 and clarified by centrifugation at 4°C, 13,000×g for 5 min. The resultant clarified mixture (~150 μl) was then loaded onto a 10-20% w/v 2.5 ml linear sucrose gradient additionally containing 0.5 ml of 50% sucrose cushion in the bottom, made with buffer 1 (25 mM HEPES-KOH pH 7.6, 100 mM KCl, 5 mM MgCl2, 0.1 mM EDTA 5 mM DTT. The gradients were prepared using freeze-thaw method^42^ in Thinwall Ultra-Clear ultracentrifuge tubes (5 ml, 13×51 mm; Beckman-Coulter). To create the 50% sucrose cushion, upon the gradients melting and stabilising overnight at 4°C, 0.5 ml of 50% sucrose in buffer 1 were injected to the bottom of the tubes with a syringe-attached glass capillary. Tubes were then ultracentrifuged in an SW 55 Ti rotor at 4°C, at 55,000 rpm, average g-force 287,980×g (k-factor 49), for 1 h 30 min. These conditions have been pre-optimised (using post-ultracentrifugation gradient absorbance trace analysis) to retain the ‘free’ (non-(poly)ribosomal) SSUs and LSUs in the top (10-20% sucrose) portion of the gradient while concentrating the (poly)ribosomal fraction in the bottom (50%) sucrose cushion without pelleting the material.

The (poly)ribosomal fraction collected at the bottom of the tubes was concentrated to 100 μl using Ultracel-10 regenerated cellulose membrane with 10 kDa molecular weight cut-off, in Amicon Ultra-0.5 ultrafiltration devices (Merck). To achieve partial removal of the sucrose, the initial concentrate was further diluted 4 times using buffer 1 and concentrated again to 200 μl (Poly)ribosomal presence in the resultant mixture was confirmed by absorbance readout of an analytical sucrose gradient ultracentrifugation run. The mixtures were stored frozen at −80°C and used as input material for the ‘total translated’ (*tt*) RNA-seq library construction, or the RNase digestion step of the TCP-seq library construction.

### RNase digestion of the fixed (poly)ribosomal complexes and separation of digested material into small ribosomal subunit (SSU), mono(ribo)somal (ribosomes, RS) and di(ribo)somal (disomes, DS) fractions

The procedure generally followed our approach as described before^40,41^ and was updated with the better-suited ultracentrifugation parameters allowing to achieve higher resolution across all collected fractions^36^. The *tt* fraction from the previous step was digested using 4.5 U of *E. coli* RNase I (Ambion) per 1 OD260 unit of the fraction for 30 minutes at 23° C. SUPERaseIn RNase inhibitor (Thermo Fisher Scientific) was immediately added to the mixture to 0.25 U/μl to inactivate RNase I, and the samples were transferred to ice. The reaction mixtures were loaded onto 12.5 ml linear 10-40% w/v sucrose gradients formed in 13 ml thinwall polypropylene tubes, 14×89 mm (Beckman-Coulter) using the freeze-thaw method^42^. Due to the different concentration of the SSU, RS and DS, fractions and the resolving power of the gradients for each fraction, different loads of the material were used to achieve optimal separation. For the SSU and RS 13-14 AU, and for DS 10-11 AU per gradient were usually taken. Minimally two gradients were used per each DS purification. The tubes with the loaded gradients were centrifuged in an SW 41 Ti rotor at 4°C, average g-force 178,305×g (k-factor 143.9), for 3 h 30 min. Absorbance profiles of the resultant sucrose gradients with the sedimentation-separated material were read at 254 nm, 1.5 ml/min, using Gradient Fractionator instrument (Brandel). Fractions corresponding to the position and mobility of the SSU, RS and DS complexes were identified in real time and isolated, their average absorbance at 260 nm recorded, and the fraction material further stored at −80°C or processed immediately.

### Construction of the TCP-seq SSU, RS and DS footprint RNA-sequencing libraries

We used an approach principally similar to the previously used by us^40,41^ and based on RNA 3’ polyadenylation, oligo(dT)-dependent reverse transcription, cDNA circularisation and PDD-based depletion of ribosomal RNA^43,44^, with several streamlining modifications and an expanded ribosomal (r)RNA depletion probe set (**Supplementary Table 1**). ~3.0 AU at 260 nm of the gradient-separated SSU, RS and DS material were used per each library.

To de-block the crosslinks and isolate the RNA, sucrose gradient fractions (350 μl) were supplemented with 40 μl of 100% stop solution (10% SDS w/v and 100 mM EDTA), Tris-HCl (pH 2 at 25°C) to 10 mM (4 μl 1 M), glycine to 10 mM (1.6 μl 2.5 M) and deionised nuclease-free water to obtain 400 μl as the final volume. Acidic Phenol:chloroform:isoamyl alcohol 125:24:1 (pH 4.0-5.0) (Merck/Sigma-Aldrich) was added followed by vigorously shaking the mixtures using vortex set to the maximum speed for 2 minutes, and continuing shaking at 65°C, 1,400 rpm for 30 min in a thermomixer (Eppendorf). Phase separation was facilitated by centrifuging the mixture at 12.000×g for 10 min at room temperature. The aqueous phases were then collected and transferred to fresh 1.5 ml Eppendorf tubes. RNA was precipitated by adding 0.1 volumes of 3 M sodium acetate (Invitrogen/Thermo Fisher Scientific), 20 μg of glycogen (Invitrogen/Thermo Fisher Scientific) and 2.5 volumes of absolute ethanol (Merck/Sigma-Aldrich). The tubes were vortexed for 1 min and incubated at −20°C for at least 2 hours. RNA was pelleted by centrifugation at 12,000×g for 30 min at room temperature. The supernatant was discarded and the pellet was washed twice with 80% v/v ethanol by centrifugation at 12,000×g for 30 min at room temperature. The RNA pellets were dried at 45°C for 10 min, the resultant dried pellet was dissolved in 20 μl of 1× HE buffer (final concentration 10 mM HEPES-KOH, pH 7.6 at 25°C, and 0.25 mM EDTA, pH 8.0 at 25°C) and RNA concentration was estimated using absorbance spectrum measurement *via* Nanodrop spectrophotometer (Thermo Fisher Scientific) and RNA quality was further assessed using RNA 6000 Pico Kit and Bioanalyzer 2100 (Agilent).

To reverse transcribe the RNA and introduce stand-specific identifiers, ~8 pmol (calculated using the average length of ~300 nt and amount of ~1μg) of the RNA per each library was taken in 20 μl of 1× HE buffer solution. The RNA solution was transferred into a low DNA binding 1.5 ml tube (Eppendorf), heated at 70°C for 2 min and immediately transferred to ice for 5 min. The RNA was then end-repaired with 20 U of 3’-phosphatase-positive bacteriophage T4 polynucleotide kinase (T4 PNK; New England Biolabs), using conditions recommended by the supplier (1× PNK buffer without ATP, 2 U/μl RNaseOUT Recombinant Ribonuclease Inhibitor (Thermo Fisher Scientific), incubation at 37°C for 2 h). The reaction was stopped and PNK inactivated by the addition of 17 μl of deionised water and heating of the reaction mixture for 20 min at 65°C. The end-repaired RNA was next 3’ polyadenylated using 0.4 U/μl of *E. coli* poly(A) polymerase (EPAP; New England Biolabs), generally according to the suppliers’ recommendations (1× EPAP buffer, 1 mM ATP, 1 mM DTT, and 0.8 U/μl RNaseOUT Recombinant Ribonuclease Inhibitor (Thermo Fisher Scientific), incubation at 37°C for 1 h). The reaction was stopped by the addition of 15 μl of 100% stop solution and the resulting polyadenylated RNA was ethanol-precipitated, washed, dried as described before, and dissolved in 25 μl of 1× HE. The RNA was next reverse transcribed using the oligo(dT) primer (5’-phosphate-GA TCG TCG GAC TGT AGA ACT CTG AAC G /9-carbon spacer/ G TGA CTG GAG TTC CTT GGC ACC CGA GAA TTC CAT TTT TTT TTT TTT TTT TTT TVN-3’) with SuperScript IV reverse transcriptase (Invitrogen/Thermo Fisher Scientific), generally as recommended by the supplier. The template RNA (up to 4 pmol) was first mixed with 20 pmol of the split adaptor primer,0.5 mM (each) dNTPs mixture, 1× SuperScript IV buffer and annealed by heating to 75°C for 3 min, then cooling to 65°C, and slow ramping (3°C/s) to 55°C. The reaction mixture was then supplemented with 5 mM DTT, 2 U/μl RNaseIn Plus (Promega) and 10 U/μl SuperScript IV Reverse Transcriptase enzyme while heated, slow-ramped to 50°C and incubated at 50°C for further 30 min.

To purify the resultant cDNA away from the excess of the RNA and the split adaptor primer, and create a strand-specific amplifiable template, the reaction mixture was snap-cooled to 37°C, supplemented with 20 U of *E. coli* exonuclease I (New England Biolabs) and incubated at 37°C for 20 min. The reaction was then stopped by the addition of 15 μl of 100% stop solution. Samples were then subjected to phenol:chloroform:isoamyl alcohol 25:24:1, pH 7.7-8.8 (Merck/Sigma-Aldrich) extraction. Aqueous phase separation and nucleic acid ethanol precipitation were performed as described earlier. The resultant purified cDNA was dissolved in 9 μl of 1× HE and liberated from RNA contaminants by heating at 85°C for 3 min, snap-chilling on ice for 5 min, supplementing with 1 μl of RNase A/T mix (2 mg/ml of RNase A and 5,000 U/ml of RNase T1, Thermo Fisher Scientific), and incubating the resulting mixtures for 20 min at 37°C.

80 pmol of empty split adaptor circularisation blocker (5’-TTN BAA AAA AAA AAA AAA AAA AAA /iSuper-dT/GG AAT TCT CGG GTG CGT GTG T/3BioTEG/-3’) and 2 μl of 10× Annealing Buffer (250 mM HEPES-KOH pH 7.6 at 25°C, 1 M NaCl and 50 mM MgCl2) were then added to a final volume of 20 μl and annealing performed by heating the mixtures at 80°C for 2 min, ramping up to 95°C for 1 min, cooling down to 50°C for 1 min and slow-ramping (3°C/min) to 40°C. The RNA-depleted cDNA was then circularised at 40°C according to the manufacturer’s instructions, by adding 4 μl of 1× CircLigase II Buffer (Epicentre/Illumina), 2 μl of 50mM of MnCl2, 8 μl of 500 mM betaine, and 1 μl of 2.5 U of CircLigase II ssDNA Ligase (Lucigen), and incubating the resultant mixtures at 40°C for 3 h. The reaction was stopped by addition of 10 μl of 100% stop solution. 1 volume of neutral phenol:chloroform:isoamyl alcohol (pH 7.7-8.8) (Sigma-Aldrich/Merck) was then added and the mixtures intensively vortexed for 2 min. Separation of the aqueous and phenol phases was facilitated by centrifugation at room temperature, 12,000×g, for 10 min. Extracted circularised cDNA in the water phase (~50 μl) was further gel-filtered using 1x HE-equilibrated Illustra MicroSpin G-25 Columns (Sigma-Aldrich/Merck) according to the manufacturer’s recommendations. To remove the empty split adaptor circularisation blocker, 23 μl (out of ~50 μl) of the gel-filtered material were mixed with 40 pmol of the block removal oligonucleotide (5’-ACA CAC GCA CCC GAG AAT TCC ATT TTT TTT TTT TTT TTT T/iSuper-dT/T VNA A/3BioTEG/-3’), and supplemented with 3 μl of 10× Annealing Buffer. The mixtures were heated at 80°C for 2 min, 95°C for 1 min, 80°C for 10 s, followed by slow-ramping to 50°C, further incubated at 50°C for 1 min and cooled to 40°C. The reaction mixtures were transferred into new 1.5 ml low DNA binding tubes (Eppendorf) containing hydrophilic streptavidin magnetic beads (equalling to 260 μl of the original bead suspension; New England Biolabs), pre-equilibrated with 1× Annealing Buffer. The tubes with reaction mixtures and beads were incubated at room temperature for 10 min with gentle tube flicking to keep the beads suspended, followed by an incubation at 35°C for 2 min, immediate separation of the beads on a magnetic rack and collection of the supernatant containing the unbound circularised cDNA.

To specifically deplete cDNA representing fragments with rRNA sequences, the circularised cDNA was supplemented with 1.9 μM of each of the depletion DNA probes (**Supplementary Table 1**) and heated at 95°C for 3 min, cooled to 75°C (3°C/min), supplemented with 1× Duplex-Specific Nuclease (DSN) buffer (Evrogen), slow-ramped (3°C/min) to 60°C and further supplemented with 0.5 U DSN enzyme while at 60°C. The reaction mixtures were then slow-ramped (3°C/min) to the hybridisation temperature of 48°C and further incubated at 48°C for 20 min. The reactions were stopped by the addition of 20 mM EDTA and 55 μl of 1×HE buffer, the mixture extracted with an equal volume of neutral phenol:chloroform:isoamyl alcohol 25:24:1 (~75 μl; pH 7.7-8.8; Merck/Sigma-Aldrich), as described earlier, and gel-purified using HE-equilibrated MicroSpin G-25 Columns (Merck/Sigma-Aldrich) according to the manufacturer’s recommendations.

To amplify the rRNA-depleted cDNA and perform library size-selection, the resultant purified PDD-treated cDNA was thermocycled with Platinum SuperFi DNA polymerase (Thermo Fisher Scientific), generally according to the manufacturer’s instructions and using custom primer pairs compatible with TrueSeq Small RNA Sample Preparation Kit (Illumina), bearing a unique tag for each library. The forward primers were 5’-CAA GCA GAA GAC GGC ATA CGA GAT XXX XXX GTG ACT GGA GTT CCT TGG CAC CCG AGA ATT CCA-3’ (in which XXX XXX represents Illumina’s indexing hexanucleotide sequences), and the reverse primer was 5’-AAT GAT ACG GCG ACC ACC GAG ATC TAC ACG TTC AGA GTT CTA CAG TCC GA-3’. The amplification reaction included the 25 μl of purified PDD-treated cDNA, 0.2 mM each of dNTPs, 5× SuperFi DNA polymerase buffer, 0.5 μM each of primers, 4.2 ng/μl extreme thermostable single-stranded DNA binding protein (New England Biolabs) and 0.02 U/μl SuperFi DNA polymerase (Thermo Fisher Scientific). Typically, 21 cycles were used, with a thermal profile 98°C for 5 min followed by 98°C for 30 s melting, annealing at 62°C for 30 s and extension was performed at 72°C for 45 s, this was followed for the first two cycles. For the third and subsequent cycles, melting was performed at 98°C for 30 s, annealing temperature was increased to 76°C for 30 s followed by extension at 72°C for 45 s; this was repeated for ~18 cycles. The final extension was performed at 72°C for 1 min. The amount of the amplified DNA samples was equalised per barcode using band intensity measurements obtained by imaging a respective native agarose gel test run with the samples loaded separately for each barcode and pre-stained with 6×GRGreen loading buffer (Excellgen). Libraries were then pooled together (typically by 4 per barcode), electrophoretically separated in a native agarose gel using 6×GRGreen loading buffer (Excellgen), and selected for the insert size of 10-250 nt by cutting out the respective region of the gel. The libraries were eluted from the gel by freezing the gel pieces at −20°C for 30 min in Freeze ‘N Squeeze DNA gel extraction spin columns (Bio-Rad) and recovering the solution by immediate centrifugation of the columns at room temperature, 13,000×g, for 3 min. The recovered DNA solution was subjected to neutral phenol:chloroform:isoamyl alcohol 25:24:1 (pH 7.7-8.8, Merck/Sigma-Aldrich) extraction and ethanol precipitation, as described earlier. Dried pellets were dissolved in ~20 μl 1× HE buffer, quality-controlled with capillary electrophoresis (using High Sensitivity DNA chips run in Agilent Bioanalyzer 2100), and directed to the high-throughput sequencing input.

### Construction of the total (*T*) and total translated (*tt*) RNA-sequencing libraries

RNA seq libraries were made using the *T* and *tt* fractions. For *T*, ~200 mg of the frozen cell grindate was subjected to reverse crosslinking followed by RNA extraction generally as described above. The frozen cell grindate was thawed and clarified by centrifugation to remove cell debris at 4°C, 13,000×g for 5 min. The resultant supernatant (~150 μl, ~3.0 AU) was then subjected to reverse crosslinking followed by RNA extraction as described above in the ‘Construction of the TCP-seq SSU, RS and DS footprint RNA-sequencing libraries’ section. For *tt*, 200 μl (~5.0 AU) of the filtered and sucrose-depleted (poly)ribosomal fraction from the ‘Separation of the fixed (poly)ribosomal complexes away from the non-translated fractions of cytosolic RNA’ section was used, and the reverse crosslinking performed as for *T*. ~1 μg of the RNA obtained from each, *T* and *tt*, was directed into making the rRNA-depleted RNA-seq libraries employing GENEWIZ services. Generally, the rRNA was depleted in the RNA fractions and VAHTS Total RNA-seq (HMR) Library Prep Kit for Illumina-compatible sequencing were used for library preparation, during which the rRNA-depleted RNA was fragmented and reverse-transcribed. First strand cDNA was synthesised using ProtoScript II Reverse Transcriptase with random primers and actinomycin D. The cDNA second strand was synthesised using Second Strand Synthesis Enzyme Mix, which included dACG-TP/dUTP. The double-stranded cDNA was Solid Phase Reversible Immobilisation (SPRI) bead-purified and then treated with End Prep Enzyme Mix to repair both ends and create dA-tailing in one reaction, followed by a T-A ligation to add amplification adaptors to both ends. Size selection of the adaptor-ligated DNA was then performed using SPRI beads, and fragments up to ~400 bp (with the approximate insert size of 100-300 bp) were recovered. The dUTP-labelled second strand was digested with Uracil-Specific Excision Reagent enzyme. Each sample was then amplified by PCR using Illumina’s P5 and P7 index-containing primers. The PCR products were cleaned up using SPRI beads and quantified by Nano quality control (Nano QC), to estimate library concentration. Next generation sequencing libraries were then multiplexed and paired-end sequenced to 150 nt read length, to obtain at least 100 M reads per each barcode.

### High-throughput sequencing and mapping of the reads

Sequencing was performed on a HiSeq 2500 (Illumina by GENEWIZ) using settings compatible with TrueSeq Small RNA Sample Preparation Kit (Illumina) and paired-end reads of 150 nt. Barcoded libraries were mixed in equimolar or otherwise desired proportion, and sequenced with 400 M reads per lane mode; additional lanes were invoked when necessary.

The 150 nt Illumina paired reads were subjected to a quality control using Trimmomatic (SLIDINGWINDOW: 7 nt; phred quality cut-off: 24) followed by adaptor trimming including the (A)_20_ tract from the reverse transcription primer. The reads containing no 3’ poly(A) or shorter than 17 nt were discarded. To assign reads to the genome a stepwise alignment strategy was performed as follows. Reads were first filtered to keep only those not mapping to any rRNA species (inferred from the locus: chromosome XII:450,000-491,000). Remaining reads were then mapped to a custom tRNA sequence set based on GtRNAdb (PMC4702915). Unmapped reads from this alignment were then aligned with spliced RNA sequences containing ‘misc_RNA’, ‘ncRNA’, ‘tRNA’ or ‘snoRNA’ primary tags in SacCer3. Finally, the remaining unmapped sequences were aligned to a genomic mRNA reference consisting of all protein-coding gene regions with an up- and downstream 1,000 bp flanking genomic sequences, allowing multiple mappings.

### Footprint end definition and mapping

After quality filtering and adaptor removal, the 3’ ends of the reads were used as reference points for the ribosome protected footprints. In cases when the 3’ end position of the trimmed read sequences was aligned immediately upstream of adenine nucleotides, it was impossible to resolve the exact location of the 3’ end of the corresponding original RNA fragment before polyadenylation. With these reads, the 3’ end position was randomly assigned with equal probability between all possible locations. Assignment of start codon and ORF regions was performed as described earlier^40^.

### Length-based footprint classification and quantification

RS and DS footprints were defined by dissection of the footprint length distribution of the RS and DS TCP-seq libraries with the range of 27-38 nt (‘short’) and 56-82 nt (‘long’). RPKM values were calculated for both paired RNA-seq fragments and ribosome-protected footprints.

### Calculation of the translation efficiency (TE)

‘Translation efficiency’ measures were calculated as the ratio of the normalised occupancy (RPKM) of TCP-seq RS fraction and the RNA-seq from the poly(ribo)some co-sedimenting (*T*; TE^*T*^) and the total cell lysate (*tt,* TE^*tt*^) pools along the ORF regions.

### Definition and calculation of the features used for the Stochastic Translation Efficiency (STE) modelling

STE measures were determined using the TCP-seq SSU, RS and DS profiles. ‘Original DS/RS’ refers to the unfiltered, unprocessed and otherwise unmodified ratio of the normalised occupancies of RS (using only short footprint; 27-38 nt) and DS (using only long footprint; 56-82 nt). 5’ UTR scanning obstruction ratio was calculated as the number of SSU footprint within the 5’ UTR divided by the number of footprint within the 5’ UTR and over the annotated start codon. Initiation efficiency was derived from the ratio of the normalised SSU footprints divided by the normalised RS signal over the ORF region. Finally, elongation efficiency was calculated as the ratio of the normalised RS signal around the start codon (the 5’ end of the footprint falls in the ±7 codons relative to the annotated start codon) divided by the average RS signal over the ORF region.

### Machine learning and modelling of STE

A machine-learned predictor for the real protein synthesis rate per mRNA per second taken from Riba *et al*. 2019^32^. Models were trained on two non-overlapping segments of the input data. The following features were derived and used: normalised SSU signal over the annotated 5’ UTR, normalised RS signal (only short, monosomal footprints), normalised DS signal (only long, disomal footprints), DS/RS ratio with and without footprint length filtering on the DS signal, 5’ UTR scanning obstruction ratio calculated as library normalised SSU signal within the 5’ UTR divided by SSU signal within the 5’ UTR and start codon-associated footprints, initiation efficiency ratio calculated as normalised RS quantified on the ORF region divided by the normalised SSU signal measured on the start codon (RS_*ORF*_/SSU_*START*_), elongation commencement ratio calculated as normalised RS quantified on the ORF region divided by normalised RS signal measured on the annotated start codon (RS_*ORF*_/RS_*START*_), and the annotated ORF length. All models were 10-fold cross-validated and tested with different train-test ratio (0.5, 0.6, 0.7 and 0.8) and with different 5’ UTR length cutoffs (10, 20 and 50 nt).

## DATA AVAILABILITY

All data from this study will be made publicly available through NCBI Gene Expression Omnibus (GEO; http://www.ncbi.nlm.nih.gov/geo/).

## CODE AVAILABILITY

In-house scripts used to generate data and figures will be made publicly available.

## ACKNOWLEDGEMENTS

The authors are grateful to the members of the Division of Genome Sciences and Cancer at The John Curtin School of Medical Research, The Australian National University. This work was supported by Australian Research Council Discovery Project grant (DP180100111 to T.P. and N.E.S), National Health and Medical Research Council Investigator Grant (GNT1175388 to N.E.S.) and Research Fellowship (APP1135928 to T.P.). The authors acknowledge The Biomolecular Resource Facility of The John Curtin School of Medical Research, The Australian National University, and the facilities of Microscopy Australia at the Centre for Advanced Microscopy, The Australian National University, a facility that is funded by the University and the Federal Government.

## AUTHOR INFORMATION

These authors contributed equally: Attila Horvath, Yoshika Janapala.

These authors jointly supervised this work: Nikolay E. Shirokikh, Thomas Preiss.

### Affiliations

Attila Horvath, Division of Genome Sciences and Cancer, The John Curtin School of Medical Research, and The Shine-Dalgarno Centre for RNA Innovation, The Australian National University, Canberra, ACT 2601, Australia

Yoshika Janapala, Division of Genome Sciences and Cancer, The John Curtin School of Medical Research, and The Shine-Dalgarno Centre for RNA Innovation, The Australian National University, Canberra, ACT 2601, Australia

Ross D. Hannan, Division of Genome Sciences and Cancer, The John Curtin School of Medical Research, and The Shine-Dalgarno Centre for RNA Innovation, The Australian National University, Canberra, ACT 2601, Australia; Department of Biochemistry and Molecular Biology, University of Melbourne, Parkville 3010, Australia; Oncogenic Signalling and Growth Control Program, Peter MacCallum Cancer Centre, Melbourne 3000, Australia; Department of Biochemistry and Molecular Biology, Monash University, Clayton 3800, Australia; School of Biomedical Sciences, University of Queensland, St Lucia 4067, Australia

Eduardo Eyras, Division of Genome Sciences and Cancer, The John Curtin School of Medical Research, and The Shine-Dalgarno Centre for RNA Innovation, The Australian National University, Canberra, ACT 2601, Australia

Thomas Preiss, Division of Genome Sciences and Cancer, The John Curtin School of Medical Research, and The Shine-Dalgarno Centre for RNA Innovation, The Australian National University, Canberra, ACT 2601, Australia; Victor Chang Cardiac Research Institute, Darlinghurst, NSW 2010, Australia

Nikolay E. Shirokikh, Division of Genome Sciences and Cancer, The John Curtin School of Medical Research, and The Shine-Dalgarno Centre for RNA Innovation, The Australian National University, Canberra, ACT 2601, Australia

### Contributions

N.E.S. and T.P. designed research, Y.J., A.H. and N.E.S. performed experiments, A.H., N.E.S., T.P., Y.J., R.D.H and E.E. analysed and interpreted the data, N.E.S. and T.P. funded research, all authors wrote the manuscript.

### Corresponding authors

Nikolay E. Shirokikh, (+61) 432847526

Thomas Preiss, (+61) 478493849

